# Fluctuation relations and fitness landscapes of growing cell populations

**DOI:** 10.1101/2020.04.10.035857

**Authors:** Arthur Genthon, David Lacoste

## Abstract

We construct a pathwise formulation of a growing population of cells, based on two different samplings of lineages within the population, namely the forward and backward samplings. We show that a general symmetry relation, called fluctuation relation relates these two samplings, independently of the model used to generate divisions and growth in the cell population. Known models of cell size control are studied with a formalism based on path integrals or on operators. We investigate some consequences of this fluctuation relation, which constrains the distributions of the number of cell divisions and leads to inequalities between the mean number of divisions and the doubling time of the population. We finally study the concept of fitness landscape, which quantifies the correlations between a phenotypic trait of interest and the number of divisions. We obtain explicit results when the trait is the age or the size, for age and size-controlled models.

## 1 Introduction

While the growth of cell populations appears deterministic, many processes occurring at the single cell level are stochastic. Among many possibilities, stochasticity at the single cell level can arise from stochasticity in the generation times [1], from stochasticity in the partition at division [2, 3], or from the stochasticity of single cell growth rates, which are usually linked to stochastic gene expression [4]. Ideally one would like be able to disentangle the various sources of stochasticity present in experimental data [5]. This would allow to understand and predict how the various sources of stochasticity affect macroscopic parameters of the cell population, such as the Malthusian population growth rate [6, 7]. Beyond this specific question, research in this field attempts to elucidate the fundamental physical constraints which control growth and divisions in cell populations.

With the advances in single cell experiments, where the growth and divisions of thousand of individual cells can be tracked, robust statistics can be acquired. New theoretical methods are needed to exploit this kind of data and to relate experiments carried out at the population level with experiments carried out at the single cell level. For instance, one would like to relate single-cell time-lapse videomicroscopy experiments of growing cell populations [8], which provide information on all the lineages in the branched tree, with experiments carried out with the mother machine configuration, which provide information on single lineages [9, 10].

Here, we develop a theoretical framework to relate observables measured at the single cell level and at the population level, building on a number of theoretical works [11, 12, 3, 13] and on our own previous work on this topic [14]. Following Nozoe et al. [15], we introduce two different ways to sample lineages, namely the forward (chronological) and backward (retrospective) samplings. We show that the statistical bias present in the backward sampling with respect to the forward sampling is captured by a general relation called fluctuation relation [3, 16, 17]. This fluctuation relation depends only on the structure of the branched tree but not on the class of dynamical models defined on it. This relation has analogies with the fluctuation relations well known in Stochastic Thermodynamics [18], as first noted in [19, 20], which we further discuss here.

Three classical models of cell size control have been proposed in the literature: the ‘sizer’ in which the division is controlled by the size of the cell, the ‘timer’ in which the division depends on the age of cell, and the ‘adder’ for which the cell divides after adding a constant volume to its birth volume [21, 22, 23, 24]. We study size models, age models and also the general case of mixed models, in which the division rate is controlled by both the size and the age of the cell. Mixed models include the three policies mentioned above, for instance the added volume appearing in the adder policy can be expressed as a function of the size and the age of the cell. We develop a framework based on fluctuation relations for these models and we explore some consequences. In section 3, we introduce a framework based either on path integrals or on operators, which characterize the symmetry contained in fluctuation relations for these models.

For these specific models and for key phenotypic variables such as the size and the age, we study in section 4 a function called fitness landscape [15], which informs whether a specific phenotypic variable affects the division rate of the cell population.

### 1.1 The backward and forward processes

Let us consider a branched tree, starting with *N*_0_ cells at time *t* = 0 and ending with *N* (*t*) cells at time *t* as shown on fig. 1. We assume that all lineages survive up to time *t*, and therefore the final number *N* (*t*) of cells correspond to the number of lineages in the tree.

**Figure 1:**
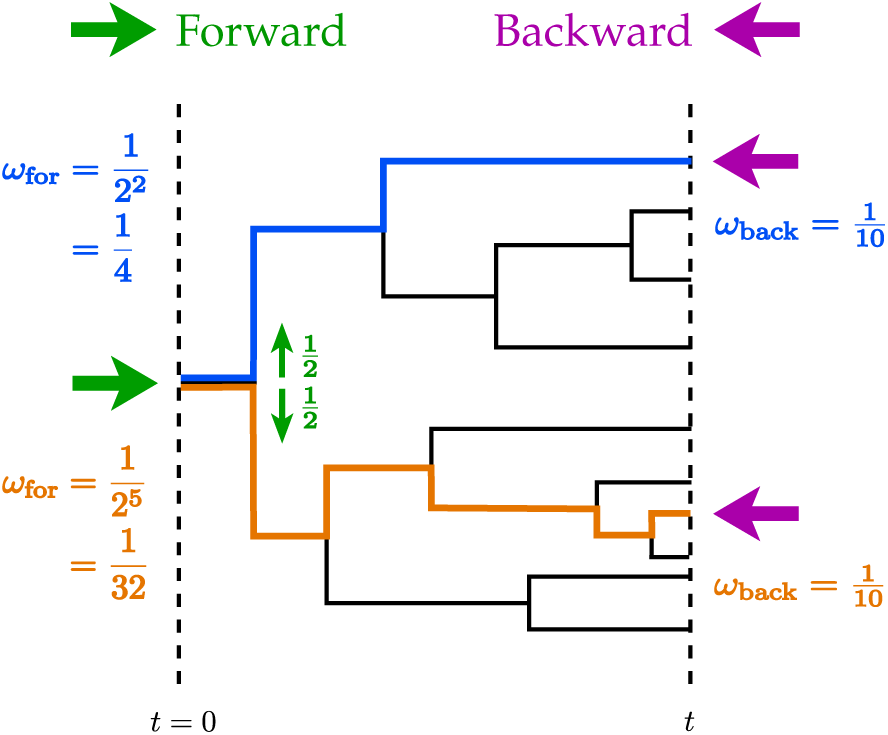
Example of a tree with *N*_0_ = 1 and *N* (*t*) = 10 lineages at time *t*. Two lineages are highlighted, the first in blue with 2 divisions and the second in orange with 5 divisions. The forward sampling is represented with the green right arrows: it starts at time *t* = 0 and goes forward in time by choosing one of the two daughters lineages at each division with probability 1/2. The backward sampling is pictured by the left purple arrows: starting from time *t* with uniform weight on the 10 lineages it goes backward in time down to time *t* = 0.

The most natural way to sample the lineages is to put uniform weight on all of them. This sampling is called the backward, (or retrospective) because at the end of the experiment one randomly chooses one lineage among the *N* (*t*) with a uniform probability and then one traces the history of the lineage backward in time from time *t* to 0, until reaching the ancestor population. The backward weight associated with a lineage *l* is defined as

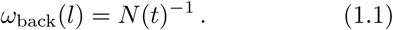

In a tree, some lineages divide more often than others, which results in an over-representation of lineages that have divided more often than the average. Therefore by choosing a lineage with uniform distribution, we are more likely to choose a lineage with more divisions than the average number of divisions in the tree.

The other way of sampling a tree is the forward (or chronological) one and consists in putting the weight

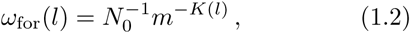

on a lineage *l* with *K*(*l*) divisions, where *m* is the number of offsprings at division. This choice of weights is called forward because one starts at time 0 by uniformly choosing one cell among the *N*_0_ initial cells, and one goes forward in time up to time *t*, by choosing one of the *m* offsprings with equal weight 1*/m* at each division. The backward and forward weights are properly normalized probabilities, defined on the *N* (*t*) lineages in the tree at time 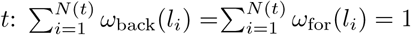.

Single lineage experiments are precisely described by a forward process since experimentally, at each division, only one of the two daughter cells is conserved while the other is eliminated (for instance flushed away in a microfluidic channel [9, 10]). In these experiments, a tree is generated but at each division only one of the two lineages is conserved, with probability 1*/*2, while the rest of the tree is eliminated. This means that single lineage observables can be measured without single lineage experiments, provided population experiments are analyzed with the correct weights on lineages.

### 1.2 Link with the population growth rate

Since the backward weight put on a lineage depends on the number of cells at time *t*, it takes into account the reproductive performance of the colony but it is unaffected by the reproductive performance of the lineage considered. On the contrary, the forward weight put on a specific lineage depends on the number of divisions of that lineage but is unaffected by the reproductive performance of other lineages in the tree. Therefore, the difference between the values of the two weights for a particular lineage informs on the difference between the reproductive performance of the lineage with respect to the colony.

We now introduce the population growth rate:

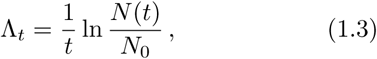

which is linked to forward weights by the relation

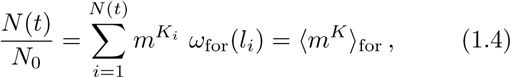

where 〈⋅〉_for_ is the average over the lineages weighted by *ω*_for_, and *K*_*i*_ = *K*(*l*_*i*_). Combining the two equations above, we obtain [25]:

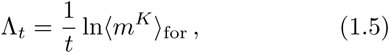

which allows an experimental estimation of the population growth rate from the knowledge of the forward statistics only.

Eq. (1.4) can also be re-written to express the bias between the forward and backward weights of the same lineage

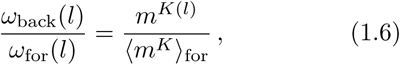

which is the reproductive performance of the lineage divided by its average in the colony with respect to *ω*_for_.

A similar relation is derived using the relation

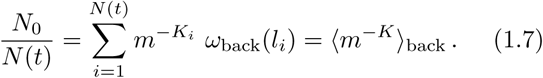

Combining eqs. (1.5) and (1.7) we obtain:

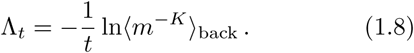

As for eq. (1.4), eq. (1.7) is reshaped to show the bias between the forward and backward weights of the same lineage:

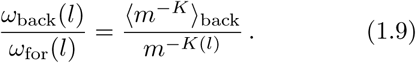

Combining eqs. (1.1) to (1.3), we obtain the fluctuation relation [14, 15]:

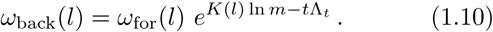

If we now introduce the probability distribution of the number of divisions for the forward sampling *p*_for_(*K*) = Σ_*l*_ *δ*(*K* − *K*(*l*))*ω*_for_(*l*) and similarly for the backward sampling, we can also recast the above relation as a fluctuation relation for the distribution of the number of divisions:

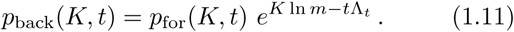

Let us now introduce the Kullback-Leibler divergence between two probability distributions *p* and *q*, which is the non-negative number:

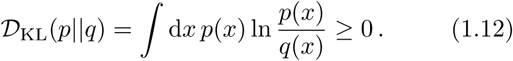

Using eq. (1.10), we obtain

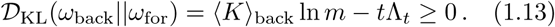

A similar inequality follows by considering *D*_KL_(*ω*_for_||*ω*_back_). Finally we obtain

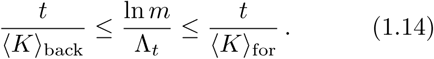

In the long time limit, 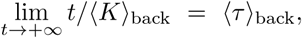 where *τ* is the inter-division time, or generation time, defined as the time between two consecutive divisions on a lineage. The same argument goes for the forward average. In the case of cell division where each cell only gives birth to two daughter cells (*m* = 2), the center term in the inequality tends to the population doubling time *T*_*d*_. Therefore, this inequality reads in the long time limit:

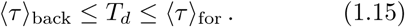

Let us now mention a minor but subtle point related to this long time limit. For a lineage with *K* divisions up to time *t*, we can write 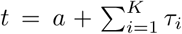, where *a* is the age of the cell at time *t* and where *τ*_*i*_ is the generation time associated with the *i*^th^ division. Then *t/K* = *τ*_*m*_ + *a/K*, where *τ*_*m*_ is the mean generation time along the lineage. For finite times, all we can deduce is *t/K* ≥ *τ*_*m*_. Therefore the left inequality of eq. (1.15) always holds

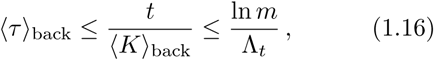

while the right inequality does not necessarily hold at finite time.

The inequalities of eq. (1.15) have been theoretically derived in [26] for age models and these authors have also experimentally verified them using experimental data. In our previous work [14], we have replotted their experimental data for clarity and we have shown theoretically that the same inequalities should also hold for size models. In fact, as the present derivation shows, the relation eq. (1.14) is very general and only depends on the branching structure of the tree, while the relation eq. (1.15) requires in addition the existence of a steady state. These inequalities and eq. (1.11) express fundamental constraints between division and growth, which should hold for any model of this type (size model, age model or mixed size-age model), irrespective of the precise form of the division rate or on the partition at cell division.

### 1.3 Stochastic thermodynamic interpretation

The results derived above have a structure similar to that found for non-equilibrium systems in Stochastic Thermodynamics [18]. Indeed, eq. (1.5) is the analog of the Jarzynski relation while eq. (1.11) is similar to the Crooks fluctuation relation, with the number of divisions *K* the analog of the work, and the population growth rate the analog of the free energy. Given that the Jarzynski or Crooks fluctuation relations have been used to infer free energies from non-equilibrium measurements, we could similarly use eq. (1.5) or eq. (1.11) to determine the population growth rate from the statistics of the number of divisions within the appropriate sampling of lineages. Relations of this type show the central role played by the population growth rate in these models. In this context, the inequalities eq. (1.14) are expressing a constraint equivalent to the second law of thermodynamics, which classically follows from the Jarzynski or Crooks fluctuation relations. Fluctuation relations also take a slightly different form when expressed at finite time or at steady state, which is indeed the case here when comparing eq. (1.14) with eq. (1.15).

A difference between the two sets of results is that the work fluctuation relations involve two specific dynamics which are related by time-reversal symmetry, whereas no dynamics is needed here to derive eq. (1.5) or eq. (1.11). We now precisely introduce dynamical variables on this tree, which will provide us with the equivalent of the non-equilibrium trajectories of Stochastic Thermodynamics.

Let us introduce *M* variables labeled (*y*_1_, *y*_2_, …, *y*_*M*_) to describe a dynamical state of the system, then a path is fully determined by the values of these variables at division, and the times of each division. We call ***y***(*t*) = (*y*_1_(*t*), *y*_2_(*t*), …, *y*_*M*_ (*t*)) a vector state at time t and 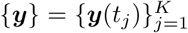 a path with *K* divisions.

The probability *P* of path {***y***} is defined as the sum over all lineages of the weights of the lineages that follow the path {***y***}:

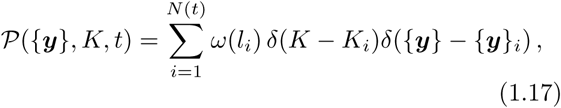

where {***y***_*i*_} is the path followed by lineage *l*_*i*_. Using the normalization of the weights *ω* on the lJineagesL, we show that 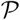 is properly normalized: 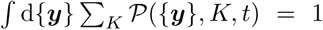. We then define the number *n*({***y***}, *K, t*) of lineages in the tree at time *t* that follow the path {***y***} with *K* divisions:

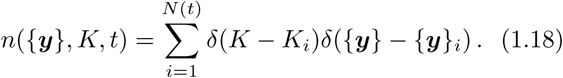

This number of lineages is normalized as ∫ d{***y***} Σ_*K*_ *n*(***y***, *K, t*) = *N* (*t*). Then, the path probability can be re-written as

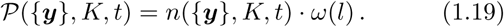

Since *n*({***y***}, *K, t*) is independent of a particular choice of lineage weighting, we obtain

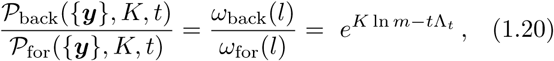

which generalizes eq. (1.11). In our previous work [14], we have derived this relation for size models with individual growth rate fluctuations (i.e. ***y*** = (*x, ν*)) but we were not aware of the weighting method introduced by [15], and for this reason, we used the term ‘tree’ to denote the backward sampling, and the term ‘lineage’ to denote the forward sampling.

## 2 Mixed age-size controlled models

### 2.1 Dynamics at the population level

The state of a cell is described by its size *x*, its age *a* and its individual growth rate ν, with ***y*** = (*x, a, ν*). The evolution of the number of cells *n*(***y***, *K, t*) in the state ***y*** at time *t*, that belong to a lineage with *K* divisions up to time *t* is governed by the equation

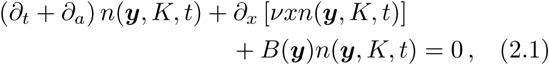

and the boundary condition

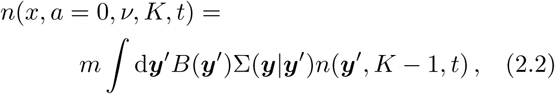

where *B*(***y***) is the division rate and Σ(***y|y**′*) is the conditional probability (also called division kernel) for a newborn cell to be in state ***y*** knowing its mother divided while in state ***y**′*, normalized as ∫ Σ(***y|y**′*)d***y*** = 1 for any ***y*′**.

### 2.2 Dynamics at the probability level

While *n*(***y***, *K, t*) in eq. (2.1) is independent of the choice of weights put on the lineages, we now turn to a description in terms of the probability *p*(***y***, *K, t*) for a cell to be in state (*y,K*) at time *t* if chosen randomly among the *N* (*t*) cells in the tree at that time. To do so, one has to choose how to weight each cell in the colony, which is equivalent to weight each lineage, since at time *t* each cell is the ending point of one lineage.

The first possibility is the backward sampling, for which each lineage is weighted uniformly. In this case, we define *p*_back_ as

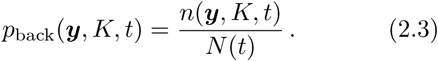

Dividing eq. (2.1) and the boundary condition eq. (2.2) by *N* (*t*) we obtain

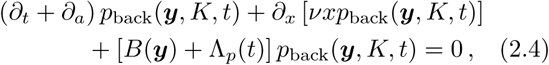

and

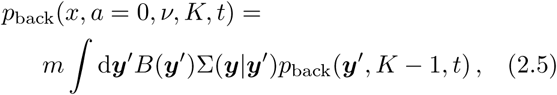

where we defined the instantaneous population growth rate as

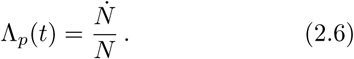

The instantaneous population growth rate and population growth rate defined in eq. (1.3) are related by:

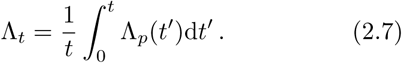

In the long-time limit, *N* grows exponentially with constant rate Λ_*p*_, and thus Λ_*t*_ = Λ_*p*_ = Λ.

The other possibility is to use the forward statistics, in which case we define the probability *p*_for_, as

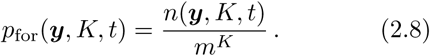

Dividing eq. (2.1) and the boundary condition eq. (2.2) by *m*^*K*^ we obtain

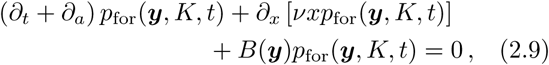

and

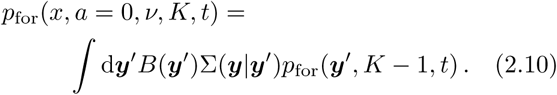

One can notice that the backward statistics is well suited to study the population, while the forward statistics reproduce the behaviour of single lineage experiments. Indeed, by taking eq. (2.4) for the population/backward probability *p*_back_, and choosing Λ_*p*_(*t*) = 0 and *m* = 1 we recover eq. (2.9). This equation is then a population equation in which we follow only one cell, so that Λ_*p*_(*t*) = 0 and *m* = 1, which we call single lineage experiment.

### 2.3 Volume conservation

Let us explain why the conservation of volume between the mother cell and the two daughter cells at division is compatible with eqs. (2.1) and (2.2). To do so, let us impose the volume conservation and show that the resulting dynamical equations to eqs. (2.1) and (2.2).

For simplicity we consider the particular case of a size-control mechanism, described in appendix A.2, with constant individual growth rate, which means that ***y*** = *x*. Our derivation would easily extend to mixed models since age does not play any role in the conservation of volume.

One can consider that when a cell of size *x*_0_ divides, the volume of one of the two daughter cells is randomly distributed according to the kernel Σ(*x*_1_|*x*_0_), and the volume of the second daughter is imposed by the conservation of volume: *x*_2_ = *x*_0_ − *x*_1_. Thus, considering the evolution of *n*(*x*_1_, *K, t*), the source term reads

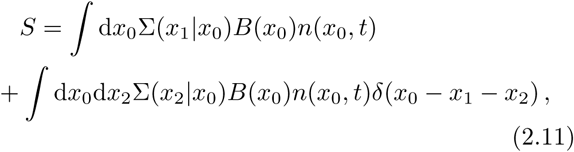

where the first term accounts for cells of size *x*_0_ randomly dividing into cells of size *x*_1_, and where the second term describes the divisions of cells of size *x*_0_ randomly dividing into cells of size *x*_2_, thus giving birth to cells of size *x*_0_ − *x*_2_ by conservation of the volume. This result can be reshaped as

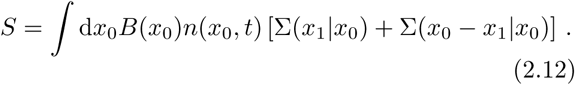

We now define a modified division kernel Σ, as 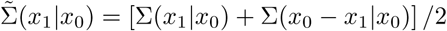, leading the source term to have the same form as in eq. (A.10), but for the modified kernel.

Moreover if Σ(*x|x′*) is symmetrical around *x* = *x′*/2, then we have the exact correspondence 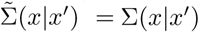.

### 2.4 Test of the fluctuation relation

We simulated the time evolution of colonies of cells, obeying eqs. (2.1) and (2.2), for age and size models in order to test the fluctuation relation. Since results are very similar -as expected- for age models, we restrict ourselves to size models. We tested two results: the fluctuation relation for the number of divisions eq. (1.11) and one of its consequences: the inequality for the mean number of divisions eq. (1.14).

All simulations for size models were conducted with the division rate *B*(*x, ν*) = *νx*^*α*^, where *α* is the strength of the control and *x* is the dimensionless size. Power law were found to be good approximations for empirical division rates *B*(*x*) as function of size [2, 27, 23], and for *B*(*a*) as function of age [27]. The factor ν, being the only time scale for size models, give *B*(*x*) its proper dimension. For age models, we chose *B*(*a, ν*) = *νf* (*aν*), where *a* is the age of the cell and with a power law dependence on age: *f* (*u*) = *u*^*α*^. The constant *ν*^*α*+1^ is redefined as *ν* since it does not play a role, so that *B*(*a, ν*) = *νa*^*α*^.

On fig. 2, the backward and forward probability distributions of the number of divisions are shown for a size model. The two distributions intersect at the number of divisions *K* = *t*Λ_*t*_/ ln 2. The inset of fig. 2 shows the logarithm of the ratio *q*(*K, t*) = *p*_back_(*K, t*)*/p*_for_(*K, t*) of the two distributions, which is as expected a straight line of slope ln 2 when plotted against the number of divisions. For convenience and for fig. 2 only, noise in the volume partition at division has been introduced, by choosing for the conditional probability Σ(*x|x′*) a uniform distribution between sizes *x* = 0 and *x* = *x′*. This has the effect of broadening the distributions *P* (*K*) with respect to the case of deterministic symmetrical volume partition.

**Figure 2:**
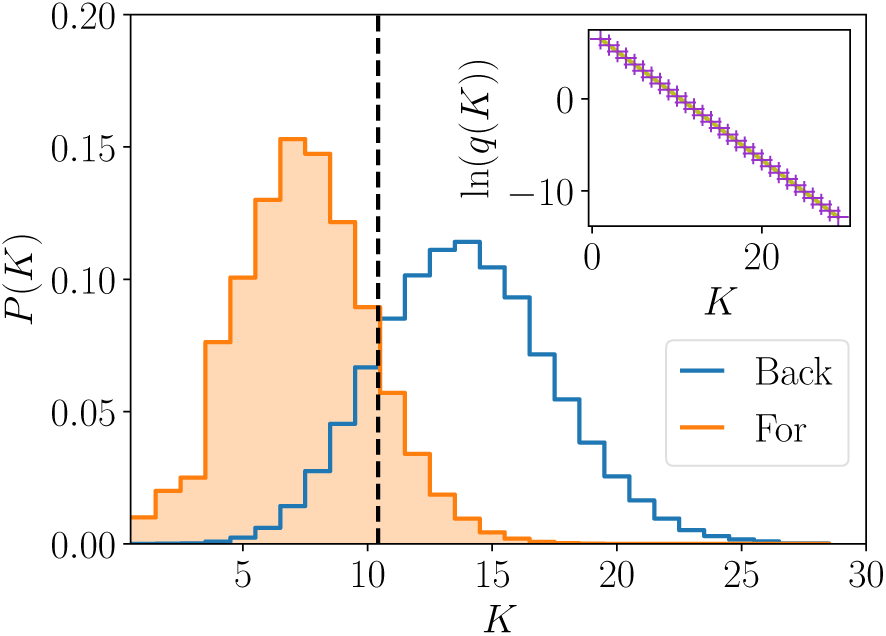
Distributions of the number of divisions in orange filled histogram for the forward statistics, and with the blue empty histogram for the backward statistics, for a size model with division rate *B*(*x*) = *νx*^*α*^, constant growth rate *ν* = 1, *α* = 2 and *t* = 7. The vertical dashed line at *K* = *t*Λ_*t*_/ln 2 is the theoretical value at which the two distributions should intersect. The inset shows the logarithm of the ratio *q*(*K*) of the forward to backward probabilities (purple crosses), and the theoretical result *t*Λ_*t*_ − *K* ln 2 (green line).

Then, we tested the inequality on the mean numbers of divisions by varying the strength of the size-control α. Results are shown on fig. 3. One one hand, we see that the less control on size, the more discrepancy between the two determinations 〈*K*〉_back_ and 〈*K*〉_for_. On the other hand, when increasing the control, the two determinations converge to the population doubling time, where no stochasticity in the number of divisions is left, and every lineage carries the same number of divisions, leading to the equality of the backward and forward statistics.

**Figure 3:**
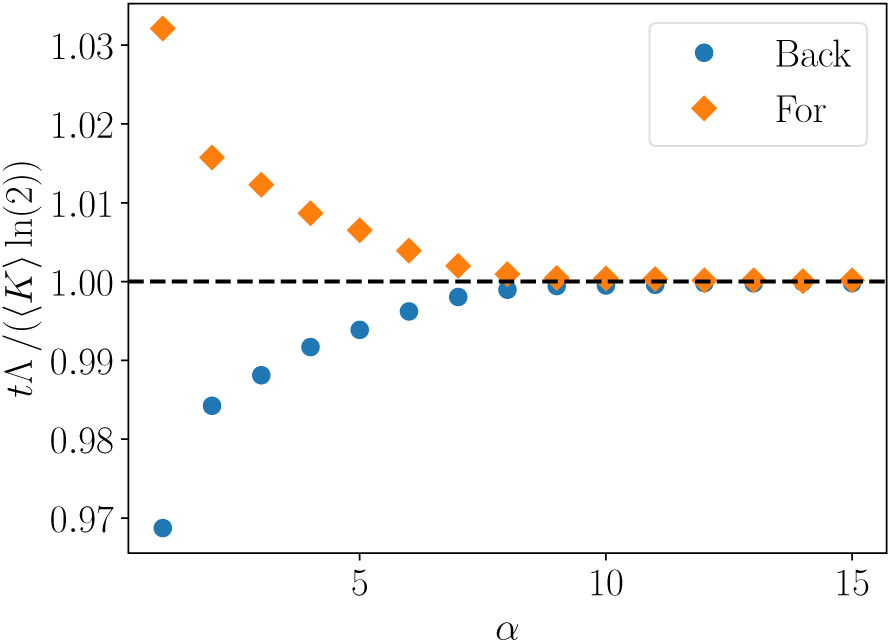
Test of the inequality on the mean number of divisions measured with the backward and forward statistics against α. The quantity *t*/〈*K*〉, re-scaled by the population doubling time *T*_*d*_ = ln 2/Λ_*t*_ is shown with orange diamonds (resp. blue circles) for the forward (resp. backward) statistics. The division rate is given by *B*(*x*) = *νx*^*α*^ where *ν* = 1, *α* is varied from 1 to 15 and *t* = 6. The volume repartition between the two daughter cells at division is symmetrical, so that Σ(*x|x′*) = δ(*x* − *x′*/2).

## 3 Path integral and operator formalisms

### 3.1 Path integral formalism

Using a path integral approach, eq. (2.1) can be solved exactly even for mixed models. A path is defined as the ensemble of values {***y***} taken by ***y*** from 0 to time *t*. Since the evolution of the size between two divisions is deterministic, the path is fully characterized by only a finite number of variables which are: the number of divisions *K*, ∀*k* ∈ [0, *K*], the size *x*_*k*_ at birth and the individual growth rate *ν*_*k*_ of the cycle, ∀*k* ∈ [0, *K* − 1] the age *a*_*k*_ at division, and the final size *x* and age *a*. Initial conditions are then given by ***y***_0_ = (*x*_0_, *a* = 0, *ν*_0_). The number of paths *n*({***y***}, *K, t*) is given by

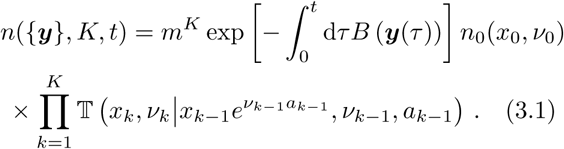

The proof of this equality and the definition of the transition matrix T are given in appendix B.

### 3.2 Recovering Wakamoto’s relation for age models

For age models, the division rate *B*(*a*) only depends on the age *a* of the cell. In this case the theoretical distribution of generation times for the forward statistics is given by

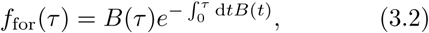

which is related to the distribution of generation times for the backward statistics *f*_back_(*τ*) by [26]:

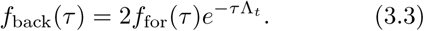

This relation can be viewed as a consequence of the general fluctuation relation at the level of path probabilities as explained below.

In the case of age models, we show in appendix B that the path probability can be reduced to

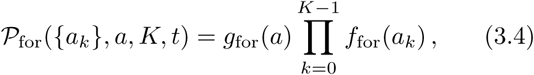

where *g*_for_(*a*) is the probability for the cell not to divide up to age *a*, between the last division at time *t* − *a* and final time *t*; and where *a*_*k*_ is the age of the cell at the (*k* + 1)^th^ division.

We define the same quantities *f*_back_(*τ*) and *g*_back_(*a*) at the backward level, obeying

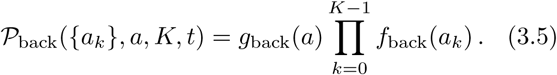

Using the fluctuation relation at the path level, namely eq. (1.20), and making a special choice for *a*_*k*_: ∀*k* ∈ [0,*K* − 1], *a*_*k*_ = *τ* = (*t* − *a*)/*K*, we obtain

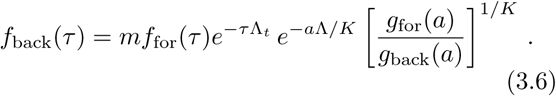

In a steady state, when *t* → ∞ and *K* → ∞, *e*^−*a*Λ/*K*^ [*g*_for_(*a*)*/g*_back_(*a*)]^1*/K*^ tends to 1 and eq. (3.6) gives back Wakamoto’s relation namely eq. (3.3), in the case *m* = 2.

### 3.3 Operator formalism

Here, we present an operator-based framework, which provides an alternate route to eq. (1.5), eq. (1.8) and eq. (1.11), that completely avoids path integrals. For simplicity, let us illustrate this formalism on the case of the size model, discussed in appendix A.2. The case of the general mixed model, which includes both age and size control can be treated along the same lines.

For the backward sampling, we define the generating function *f*_back_(***y***, *λ*, *t*):

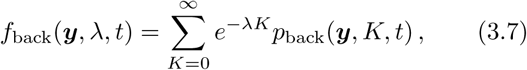

We then multiply eq. (A.12) by *e*^−*λK*^ and sum over *K* to obtain

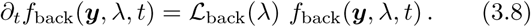

The linear operator 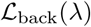, acting on *f*_back_(***y***, *λ*, *t*), is defined on a test function *f* as

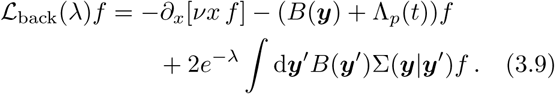

Although this operator explicitly depends on the state ***y***, we choose not to write this dependency explicitly to ease the reading.

By the same method, we obtain the operator at the forward level:

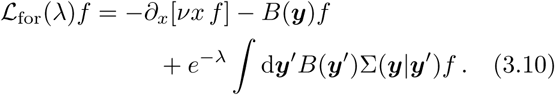

By direct comparison, we obtain the fluctuation relation at the level of operators

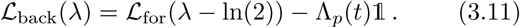

This equality between two operators implies relations between the eigenvalues and the eigenvectors as well. Let us call *χ*_back_(***y***, *λ*) (resp. *χ*_for_(***y***, *λ*)) an eigen-value of the operator 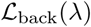 (resp. 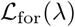), and 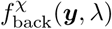 (resp. 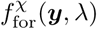) the associated eigenvector. Then the fluctuation relation at the operator level eq. (3.11) gives

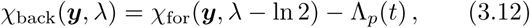

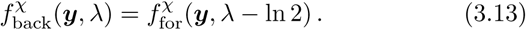

When solving eq. (3.8), the long-time behavior of *f*_back_(***y***, *λ*, *t*) is controlled by the largest eigenvalue of 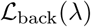, which we call *µ*_back_(***y***, *λ*), and reads

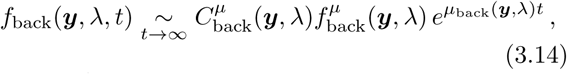

where 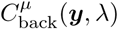 is the constant coefficient of the eigenvector 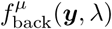 associated with the largest eigenvalue, in the decomposition of the initial condition *f*_back_(***y***, *λ*, *t* = 0) on the set of the eigenvectors of 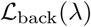.

We investigate the particular case *λ* = 0. On the one hand, using the definition eq. (3.7) we obtain

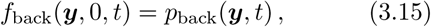

and thus the normalization of the probability gives

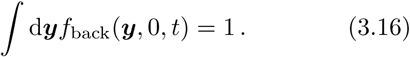

On the other hand, integrating the long time behaviour eq. (3.14) over ***y***, we get

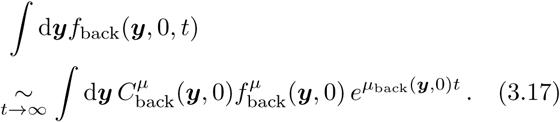

The only solution to satisfy both conditions eqs. (3.16) and (3.17) is

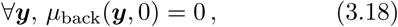

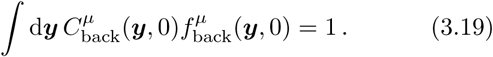

Since the largest backward eigenvalue is independent of the state ***y*** for *λ* = 0, we define the state-independent backward eigenvalue 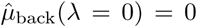. Reporting 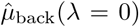 in eq. (3.12), the right hand side of the equation has to be independent of the state ***y*** as well, so we define the state-dependent forward eigenvalue: 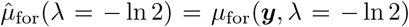 for any ***y***. Finally, eq. (3.12) gives

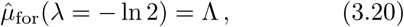

where Λ is the steady-state value of the population growth rate Λ_*p*_(*t*).

We proved that the steady-state population growth rate Λ is the largest eigenvalue of the forward (resp. backward) operator 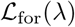 (resp. 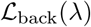) for the specific choice *λ* = −ln 2 (resp. *λ* = 0).

This choice of *λ* can be understood as follows. Let us consider the steady-state version of equation eq. (3.8) for *λ* = 0:

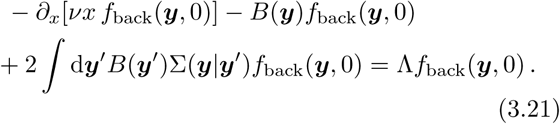

This equation, together with the normalization condition of *f*_back_(***y***, 0) that we already noted in eq. (3.16), form an eigenproblem whose unique eigenvalue is called the Malthus parameter [7], which is indeed equal to the steady-state population growth rate. Finally, the operator acting on the left hand side of eq. (3.21) differs from 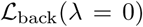, only by a term −Λ1, which gives back the value 0 for the eigenvalue of operator 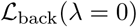.

We now use the long-time behaviour and the computed value of the largest forward eigenvalue to propose a second derivation of the link between the population growth rate and the forward statistics for the number of divisions eq. (1.5).

On the one hand, using the definition of the generation function we get

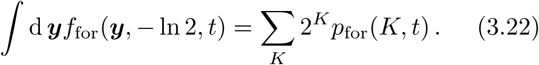

On the other hand,

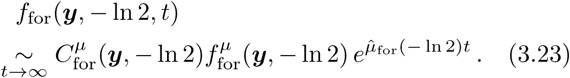

Using eqs. (3.13) and (3.19) and the fact that the initial condition is the same for both backward and forward samplings: *f*_for_(***y***, *λ*, *t* = 0) = *f*_back_(***y***, *λ*, *t* = 0) for any ***y*** and *λ*, we prove that

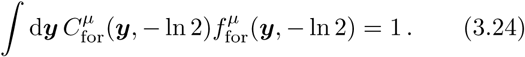

Finally, integrating eq. (3.23) over ***y*** and combining eqs. (3.20), (3.22) and (3.24), we recover

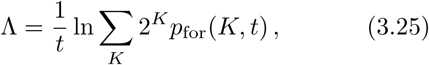

which is eq. (1.5) for *m* = 2.

## 4 Phenotypic fitness landscapes

The fitness of a phenotypic trait *s* is a measure of the reproductive success of individuals carrying it. It is usually defined as the number of offsprings of one individual with a given value of the trait and is quite difficult to evaluate. Nozoe et al. suggested that one way to measure it could be to compare the chronological and retrospective marginal probabilities [15] and accordingly defined it as:

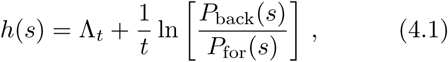

so that

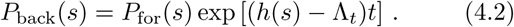

This has again the form of a fluctuation relation similar to eq. (1.11), except for the replacement of the factor *K* ln 2*/t* by the function *h*(*s*). This suggests that the fitness landscape *h*(*s*) plays a role similar to that of an effective division rate, which depends on the trait *s*. In line with this interpretation, in the particular case where *s* = *K*, eq. (1.11) leads to 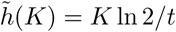, where the fitness landscape for trait *K* is called the lineage fitness and is written 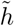. Indeed, in a branched tree, lineages with a large number of divisions *K* are exponentially over-represented in the population as compared to lineages with small numbers of divisions. This means that lineages with large *K* have a larger fitness than the ones with a small *K*, which is coherent with 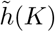 being an increasing function of *K*.

Nozoe et al. also introduced a selection pressure, which measures how the population growth rate changes when a trait *s* changes and which is defined as [15]

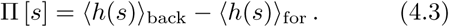

This may be written as

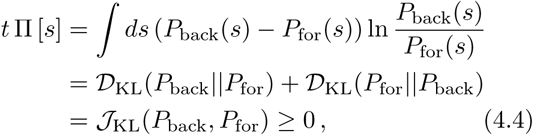

where in the last step, we introduced the Jeffreys divergence. Note that when Π [*s*] is large, the trait *s* is strongly correlated with the lineage fitness.

In the following, we rewrite the definition of *h*(*s*) in a slightly different way using

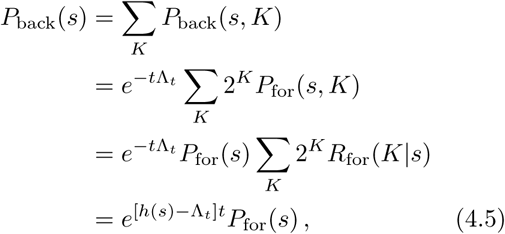

where we have introduced the probability of the number of division events conditioned on trait *s* at the forward level, *R*_for_(*K|s*). Lastly, the fitness landscape reads

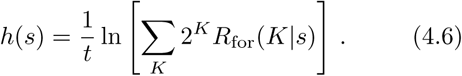

An increasing or decreasing fitness landscape means a positive or negative correlation of the trait value with the capacity to divide, whereas a constant fitness landscape means that the trait is not correlated with the number of divisions. Indeed, if we consider a trait *s* which does not affect the number *K* of divisions, then 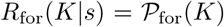 and eq. (4.6) reads 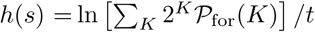, which is equal to Λ_*-t*_ according to eq. (1.5). In that case, we find that the backward and forward probabilities for that trait *s* are equal.

In the next sections, we evaluate the relevance of the key variables from our model, namely the size and the age by evaluating their fitness landscape in size and age models.

## 4.1 Size models

We start with a case where the fitness landscape is fully solvable namely a size model with no individual growth rate fluctuations and with symmetric division. These hypotheses may apply to E. Coli, which is known to show small variability in single cell growth rates, and to divide approximately symmetrically. Let us consider a colony starting with one ancestor cell of size *x*_0_. Then, the available sizes at time *t* are discrete and given by *x* = *x*_0_ exp[ν*t*]/2^*K*^ where *K* is the number of divisions undergone by the cell. Therefore a particular size *x* can be reached only if there is an integer *K* satisfying this relation, and this integer is unique, leading to

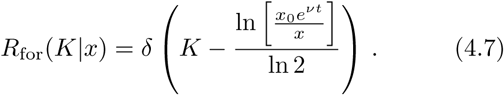

Using this relation in eq. (4.6), one finds

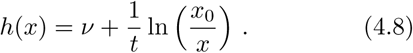

This result was tested numerically and illustrated in fig. 4.

**Figure 4:**
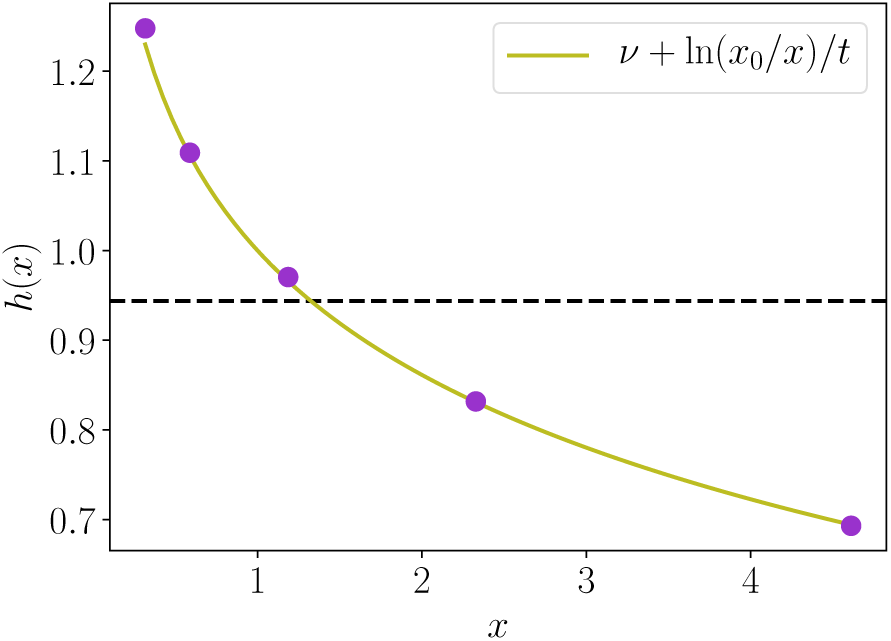
Size fitness landscape for size models with *B*(*x*) = *νx*^*α*^, constant *ν* = 1, *α* = 1, *t* = 5, an initial cell of size *x*_0_ = 1 and symmetrical division. The simulated purple dots are positioned at discrete values of *x* and the green curve is the theoretical prediction. The black horizontal dashed line represents the population growth rate Λ.

The fitness landscape of the size is a decreasing function, which implies that cells with a small size are over-represented in the population. This is coherent with the over-representation of cells that divided a lot, since these cells are more likely to be small due to the numerous divisions. Reporting this result in eq. (4.5), we obtain a fluctuation relation for the size

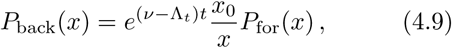

which in the long time limit becomes

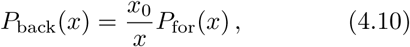

where we used the property that in a steady state, the population growth rate and the individual growth rate are equal when there is no individual growth rate variability.

In some setups, experiments do not start with a unique ancestor cell but with *N*_0_ > 1 initial cells, with possibly heterogeneous sizes. We describe this heterogeneity by the average initial size 〈*x*_0_〉 and the standard deviation 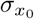. In this case, accessible sizes are still discrete but depend on both the number of divisions and the initial cell that started the lineage, and are expressed 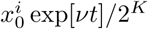, where *K* takes integer values from 0 to ∞ and where 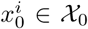, with 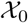 the set of initial sizes. Consequently, a final size *x* can possibly be reached by different couples 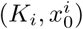.

In order to go further, we now introduce explicitly the initial sizes 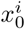 in eq. (4.6) as

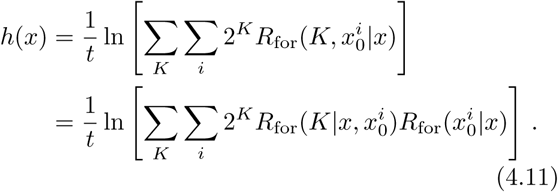

When conditioning on the initial size 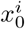, there is only one possible number of divisions *K* to reach size *x*, so that 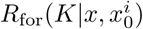 obeys an equation similar to eq. (4.7):

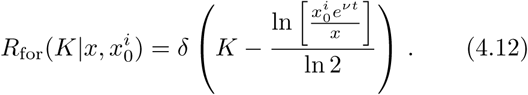

Let us examine two limit cases: (i) small variability in the initial sizes and (ii) large variability in the initial sizes. Case (i) is characterized by a small number *N*_0_ of initial cells and a small coefficient of variation 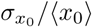. In this case, it is realistic to say that a final size *x* can only be reached by one couple (*K∗, x∗*), because the sets of accessible sizes generated by each initial cell do not overlap. Therefore, 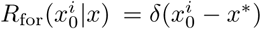 and so for any final size *x*, only one initial size *x*\* survives in the sum, so that eq. (4.11) reads

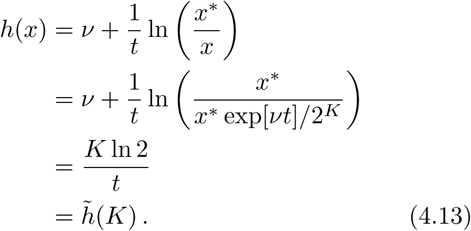

We learn from this formula that cells that come from lineages with the same number of divisions *K* have the same fitness landscape value *h*(*x*) for the size, regardless of the size *x∗* of the initial cell of their lineages, and this value is the lineage fitness 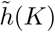. Thus, available values for *h*(*x*) are quantified by *K* and form plateaus, where points representing cells coming from different ancestors but with the same number of divisions accumulate, as shown in fig. 5a.

**Figure 5:**
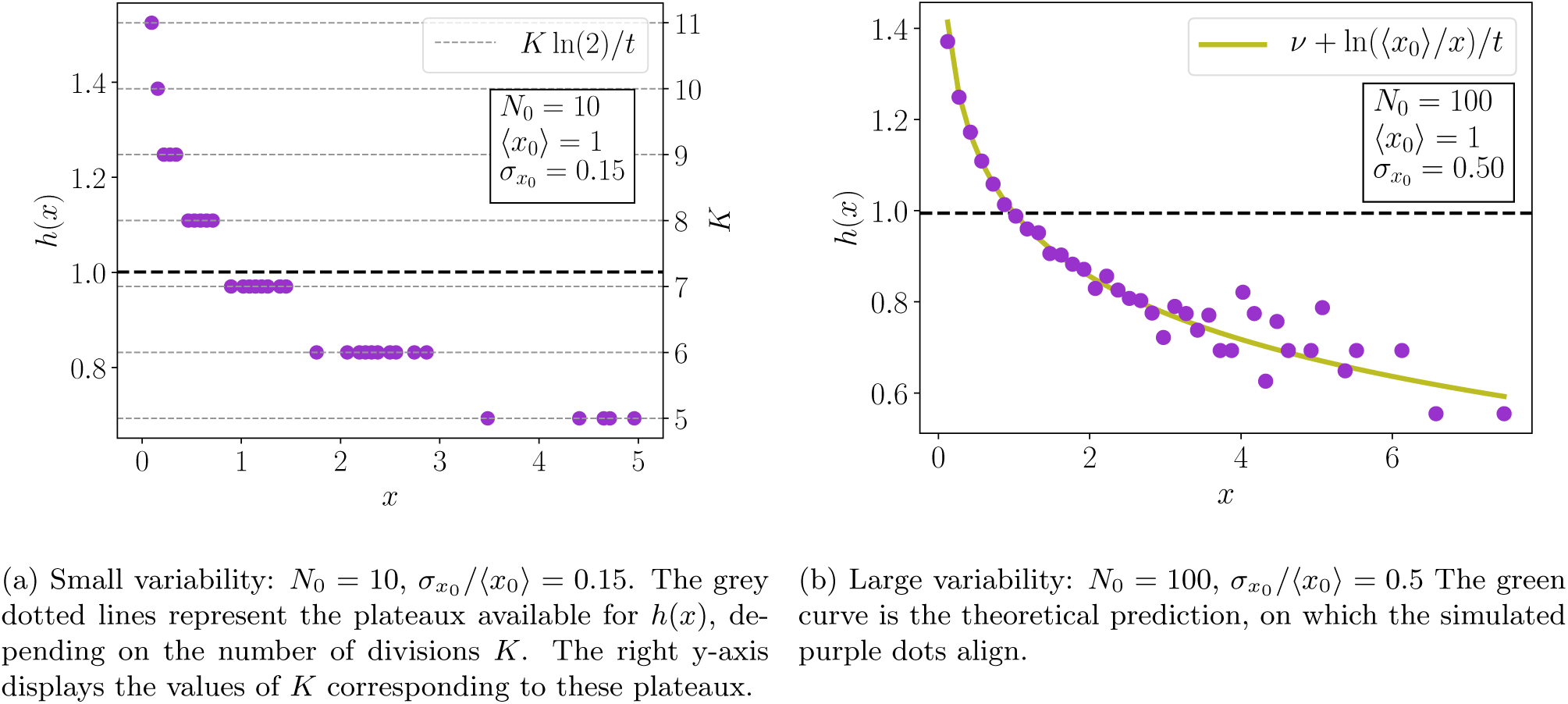
Size fitness landscapes for size models with *B*(*x*) = *νx*^*α*^, constant *ν* = 1, *α* = 1, *t* = 5, symmetrical division and *N*_0_ initial cells following a Gaussian distribution of sizes 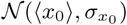. In both plots, the black horizontal dashed line represents the population growth rate Λ.

Case (ii) is characterized by a large number *N*_0_ of initial cells and a large coefficient of variation 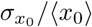. Unlike in case (i), the sets of accessible sizes generated by each initial cell have many overlaps, so that a final size *x* can be reached by many different couples 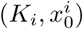. We make the hypothesis that a final *x* can be reached by any initial cell with uniform probability, so that 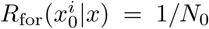. Therefore, eq. (4.11) becomes

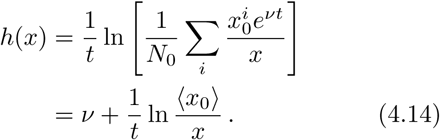

This behavior was tested numerically and the result plotted on fig. 5b confirms that the plateaus in case (i) are replaced by a smooth curve depending on the mean initial size.

We observe the same effect, namely the loss of the plateaus, when introducing fluctuations in individual growth rates.

### 4.2 Age Models

#### 4.2.1 Constant individual growth rate

We consider the case where the individual growth rate is constant and equal to *ν*. In steady-state, the forward age distribution eq. (A.8) reads (see appendix A.1 for details)

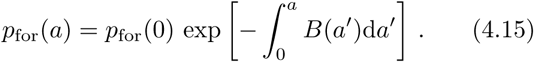

To find the integration constant *p*_for_(0), we use the normalization of probability *p*_for_:

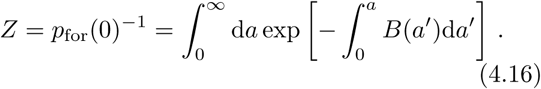

Similarly, the steady-state backward distribution of ages eq. (A.9) reads

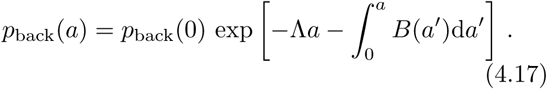

In this case, the integration constant *p*_back_(0) can be expressed both using the normalization of *p*_back_(*a*), as done for the forward case, or using eq. (A.7) which leads to *p*_back_(0) = 2Λ.

Therefore, the ratio of the age distributions using the backward and forward statistics reads

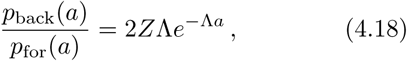

where *Z* is defined in eq. (4.16) and only depends on the division rate *B*(*a*). This relation has a similar form as the relation derived by Hashimoto et al. [26] for the distributions of generation times eq. (3.3), except for the extra age-independent factor *Z*Λ. Finally, the fitness landscape reads

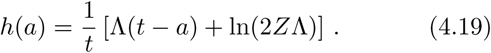

For the same reason as for *h*(*x*) in size models, *h*(*a*) in age models is a decreasing function of *a* because lineages that divided a lot are over-represented in the population and are therefore more likely to contain young cells at time *t*.

The initial condition does not play any role in this derivation, therefore, unlike size models, the results obtained are unchanged for any number *N*_0_ of initial cells with heterogeneous initial ages.

The above calculation is general because we did not put any constraint on *B*(*a*). Let us now go into more details by choosing a power law for the division rate: *B*(*a*) = *νa*^*α*^. In this case, the integral of eq. (4.16) is solvable and gives

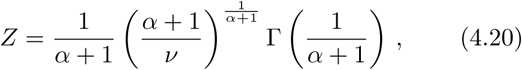

where the Gamma function is defined as

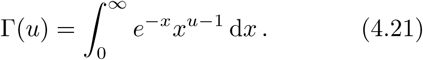

Results are plotted on fig. 6, which shows that theoretical predictions for the backward and forward age distributions are in good agreement with the numerical histograms. The inset plot shows the age fitness landscape, which follows the linear behavior predicted by eq. (4.19).

**Figure 6:**
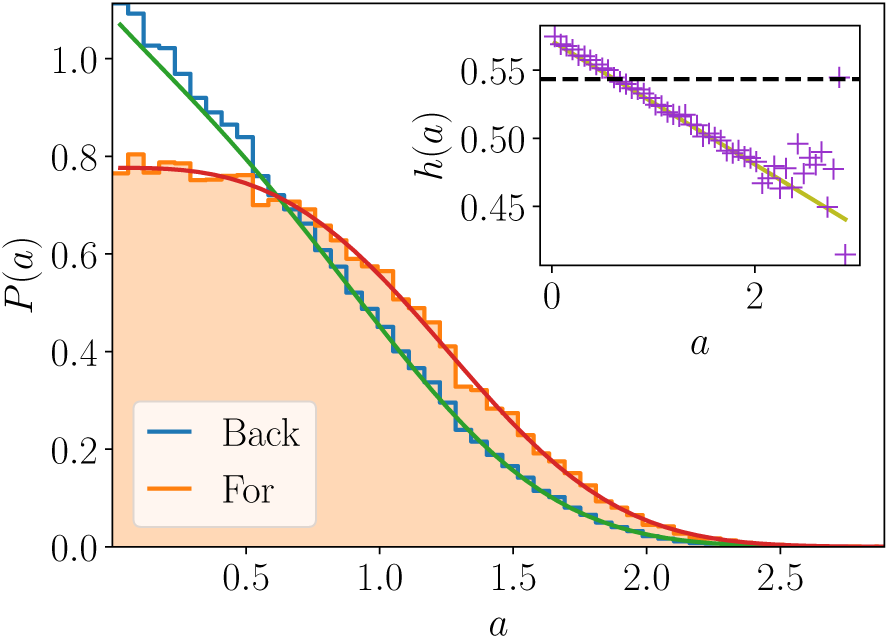
Age distributions with the forward statistics (orange filled histogram) and with the backward statistics (blue unfilled histogram), and the corresponding theoretical curves in red and green. The inset plot shows the age fitness landscape (purple crosses) and the theoretical linear law (green). The horizontal black dashed-line represents the population growth rate Λ. Simulations were conducted with *B*(*a*) = *νa*^*α*^, constant *ν* = 1, *α* = 2, *t* = 12 and *N*_0_ = 1.

Let us examine the particular case of uncontrolled models, for which the division rate is constant: *B* = *ν*. This corresponds to the case *α* = 0 in the power law analysis conducted above. Replacing *α* by 0 in eq. (4.20) leads to *Z* = 1/*ν*; moreover in steady state Λ = *ν*, so that

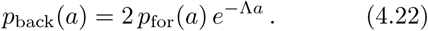

The extra factor *Z*Λ cancels and we find the same relation between the backward and forward age distributions as Wakamoto’s one between generation times distributions. Moreover, the distributions themselves are greatly simplified and read

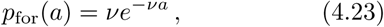

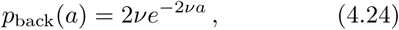

which shows that in this special case the age distributions are themselves identical with the generation time distributions.

#### 4.2.2 Fluctuating individual growth rates

We now introduce the possibility for the individual growth rate *ν* to be randomly re-defined at each division, so that the sate of a cell is now determined by two traits: ***y*** = (*a, ν*).

In this case, the steady-state joint distributions of ages and individual growth rates are given by eqs. (A.8) and (A.9). To obtain the steady-state age distributions one needs to integrate these joint distributions on individual growth rates:

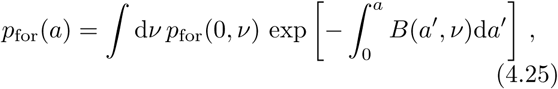

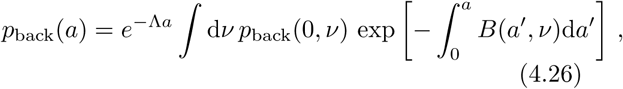

where *p*_for_(0, *ν*) and *p*_back_(0, *ν*) are given by the boundary terms eqs. (A.4) and (A.6):

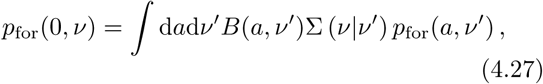

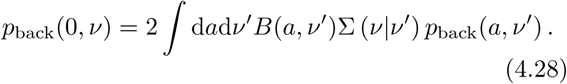

In the absence of mother-daughter correlations for the individual growth rate, then 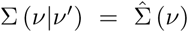, which implies that *p*_for_(0, *ν*) and *p*_back_(0, *ν*) have the same dependency in *ν*:

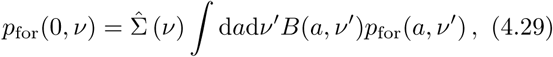

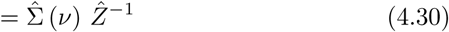

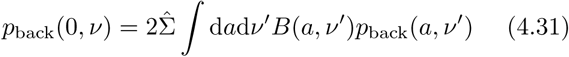

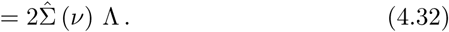

Finally, the fluctuation relation for the age reads

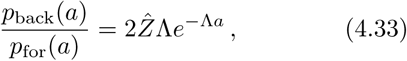

which is the equivalent of eq. (4.18) for fluctuating growth rates without mother-daughter correlations. Therefore, the age fitness landscape features the same linear dependency in age with a slope Λ as in the case of constant individual growth rate.

In the general case with mother-daughter correlations, this statement is not necessarily true though. Indeed, *p*_for_(0, *ν*) and *p*_back_(0, *ν*) do not have in general the same dependency in *ν* and therefore the integrals on *ν* in eqs. (4.25) and (4.26) do not have the same dependency in *a*, thus adding an extra age-dependent term in the age fitness landscape.

Consequently, looking at the slope of the age fitness landscape informs on the presence of mother-daughter correlations. We checked numerically that the age fitness landscape without mother-daughter correlations aligns with the theoretical prediction of slope Λ; while the same function with mother-daughter correlations presents a non-linear age dependency.

## 5 Discussion

We have studied two different methods to sample lineages in a branched tree: one sampling called backward or retrospective presents a statistical biais with respect to the forward or chronological sampling, an observation which is important to relate experiments carried out at the population level with the ones carried out at the single lineage level. This statistical bias can be rationalized by a set of fluctuation relations, which relate probability distributions in the two ensembles and which are similar to fluctuation relations known in Stochastic Thermodynamics. This analogy suggests new methods to infer the population growth rate based on statistics sampled only in the forward or backward direction. Distributions of the number of divisions are also constrained by these fluctuation relations, which can also be expressed at the level of path probabilities or at the operator level. Interesting inequalities between the mean number of divisions or the mean generation times follow from these fluctuation relations. One inequality concerning the mean generation time has already been verified experimentally by Hashimoto et al. [26] for various strains of E Coli. It would be interesting to perform more experimental investigations of this kind for other cell types or under different conditions, since such tests are very important to test our framework.

By measuring the difference in the two samplings, namely the forward and the backward one, for a specific trait, one can detect whether that particular trait is or not correlated with the division rate. The fitness landscape, related to this difference, has been introduced by Nozoe et al. [15]. While these authors have applied that concept to variables which are not reset or redistributed at division in their work, in the present paper, we used the concept of fitness landscape for variables like the size and the age, which precisely undergo a reset at division in size and age models. We derived expressions for these fitness landscapes, which agree with the statistical biais which we expect when measuring size or age distributions in cell populations. In addition, we also find that the precise form of the age fitness function appears to inform whether or not mother-daughter correlations are present in age models.

In the future, it would be interesting to further investigate models with mother-daughter correlations or with correlated single cell growth rates. It would be also valuable to extend the calculations of fitness landscapes to include other important phenotypic state variables besides size or age, as done in [28]. We hope that our work has contributed to clarifying the connection between single lineage and population statistics and to understanding the fundamental constraints which cell growth and division must obey.

## Acknowledgments

The authors acknowledge R. García-García for a previous collaboration, which made possible the present work. We would also like to thank L. Robert, P. Gaspard and J. Unterberger for stimulating discussions.

## Appendix A Age and size controlled models

### A.1 Age models

For models where the division is only controlled by the age *a* of the cell and its growth rate *ν* (so that ***y*** = (*a, ν*)), through a division rate *B*(*a, ν*), eq. (2.1) and eq. (2.2) simplify as

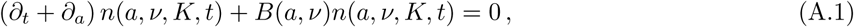

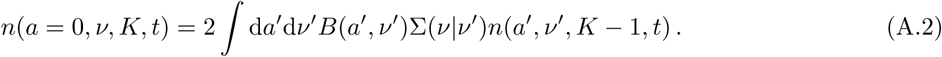

As discussed in section 2.2, the dynamics can also be expressed at the level of probabilities instead of populations. Then, the equation and boundary term for the forward probability, summed over *K*, read

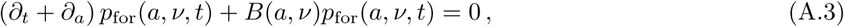

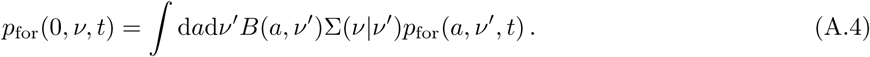

Similarly, for the backward probability, we obtain

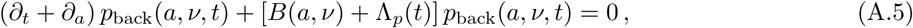

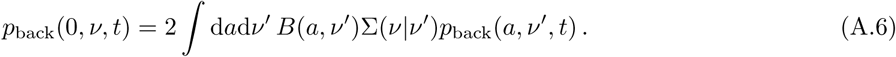

We can relate the instantaneous population growth rate Λ_*p*_(*t*) to the backward averaged division rate by integrating eq. (A.5) over *a* and *ν* and using the boundary condition eq. (A.6) and the normalization of *p*_back_:

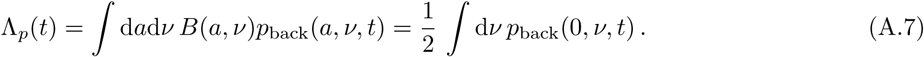

In steady-state, eqs. (A.3) and (A.5) can be solved to give the joint distribution of age and individual growth rate:

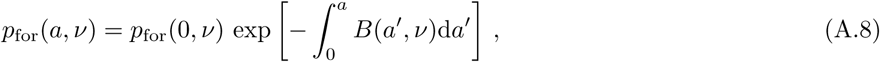

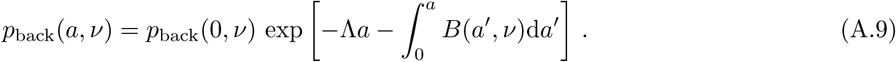

### A.2 Size models

For models where the division is only controlled by the size *x* of the cell and its growth rate *ν* (so that ***y*** = (*x, ν*)), through a division rate *B*(*x, ν*), eq. (2.1) and the boundary condition eq. (2.2) merge as

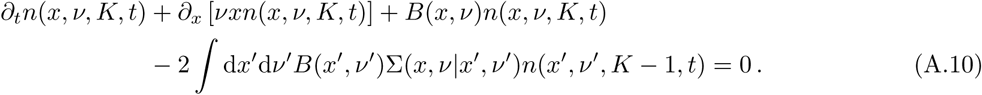

As for age models, the equation for the forward probability reads

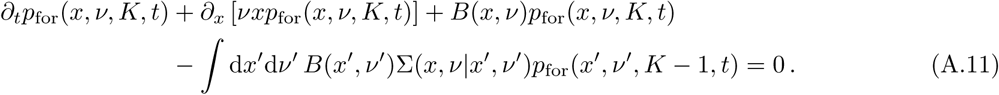

Similarly, for the backward probability, we obtain

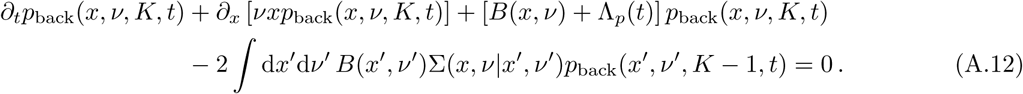

Integrating eq. (A.12) over *x* and *ν*, summing over *K* and using the normalization of *p*_back_ we obtain

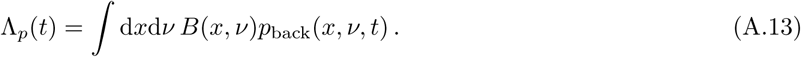

## Appendix B Path integral solution

### B.1 General solution

We start by giving a recursive solution to eq. (2.1)

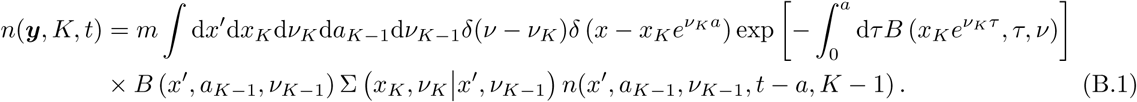

One can compute the cell number iteratively until reaching *K* = 0, which represents cells that have not divided up to time *t*. This boundary term reads

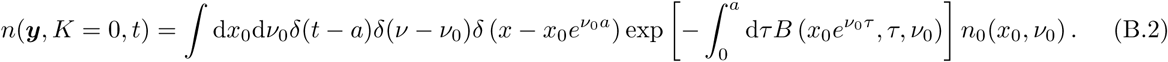

The final result then reads

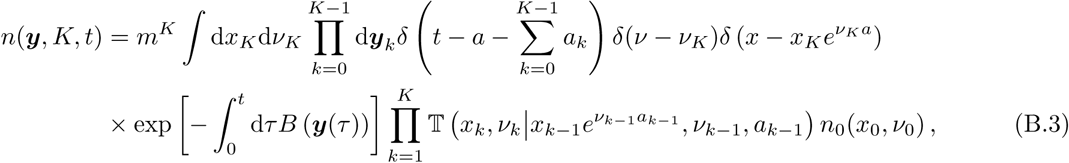

where trajectories ***y***(*τ*) explicitly appearing in the exponential are given by *ν*(*τ*) = *ν*_*k*_, *x*(*τ*) = *x*_*k*_ exp(*ν*_*k*_(*τ* − *t*_*k*_)) and *a*(*τ*) = *τ* − *t*_*k*_ for *τ* ∈ [*t*_*k*_, *t*_*k*+1_[. The *t*_*k*_ are the division times, given by 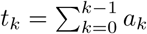 for *k* ∈ [1, *K* − 1] and *t*_0_ = 0, *t*_*K*+1_ = *t*. In addition, the transition matrix 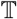 is defined as 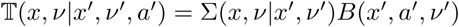.

Lastly, we can define the number of paths *n*({***y***}, *K, t*) as the number of lineages in the tree that followed the path {***y***} with *K* divisions:

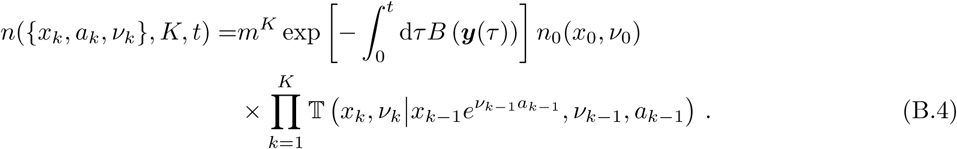

### B.2 Age models

In the case of age models with constant individual growth rate *ν*, then ***y*** = *y* = *a*, and Σ(*a|a′*) = *δ*(*a*) because new cells always appear at age 0. Therefore eq. (B.3) reads

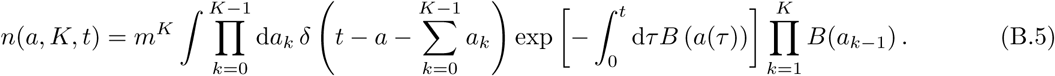

The number *n*(*{a*_*k*_}, *K, t*) of paths then reads

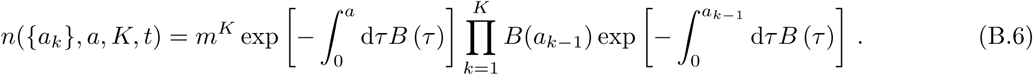

Using the definition of the forward path probability 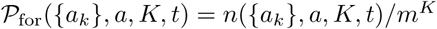 we obtain

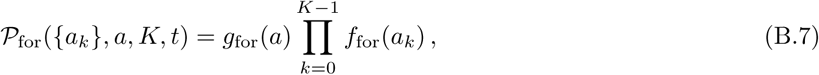

where 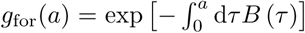 is the probability for the cell not to divide from age 0 at time *t* − *a* to age *a* at finale time *t*; and where *f*_for_ is the forward generation time distribution

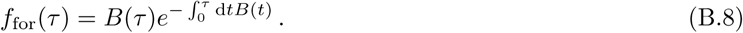

We define the equivalent quantities *f*_back_(*τ*) and *g*_back_(*a*) at the backward level:

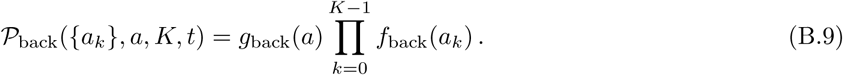

Using the fluctuation relation at the path level, namely eq. (1.20), and making the choice *∀k* ∈ [0, *K* − 1], *a*_*k*_ = *τ* = (*t* − *a*)*/K*, we obtain

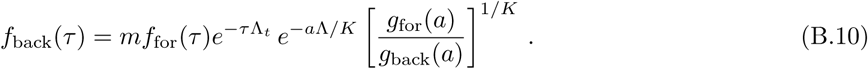

In a steady state, when *t* → ∞ and *K* → ∞, then *e*^−*a*Λ/*K*^[*g*_for_(*a*)*/g*_back_(*a*)]^1*/K*^ → 1 so the steady-state backward distribution of generation times is given by

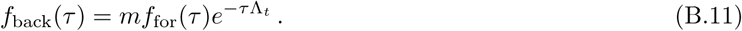

